# A transformer model for *de novo* sequencing of data-independent acquisition mass spectrometry data

**DOI:** 10.1101/2024.06.03.597251

**Authors:** Justin Sanders, Bo Wen, Paul Rudnick, Rich Johnson, Christine C. Wu, Sewoong Oh, Michael J. MacCoss, William Stafford Noble

**Affiliations:** Paul G. Allen School of Computer Science and Engineering, University of Washington; Department of Genome Sciences, University of Washington; Spectragen Informatics

## Abstract

A core computational challenge in the analysis of mass spectrometry data is the *de novo* sequencing problem, in which the generating amino acid sequence is inferred directly from an observed fragmentation spectrum without the use of a sequence database. Recently, deep learning models have made significant advances in *de novo* sequencing by learning from massive datasets of high-confidence labeled mass spectra. However, these methods are primarily designed for data-dependent acquisition (DDA) experiments. Over the past decade, the field of mass spectrometry has been moving toward using data-independent acquisition (DIA) protocols for the analysis of complex proteomic samples due to their superior specificity and reproducibility. Hence, we present a new *de novo* sequencing model called Cascadia, which uses a transformer architecture to handle the more complex data generated by DIA protocols. In comparisons with existing approaches for *de novo* sequencing of DIA data, Cascadia achieves state-of-the-art performance across a range of instruments and experimental protocols. Additionally, we demonstrate Cascadia’s ability to accurately discover *de novo* coding variants and peptides from the variable region of antibodies.

## 1 Introduction

In proteomics applications of tandem mass spectrometry, one of the core analysis challenges is the *de novo* sequencing problem: given an observed fragmentation spectrum, can we infer the amino acid sequence of the peptide responsible for generating that spectrum? Recently, significant advances have been made in *de novo* sequencing by training machine learning models that learn from a training set of labeled spectra [1]. Many of these methods [2–10] employ a type of deep neural network architecture known as a transformer [11], which was initially developed in the field of natural language processing. Accordingly, transformer models treat the *de novo* sequencing problem as a sequence-to-sequence translation task, where the spectrum is represented as a sequence of peaks and the output is a sequence of amino acids.

Over the past decade, the field of mass spectrometry has been gradually migrating to an alternative type of data generation scheme that is not compatible with most existing *de novo* sequencing tools. Traditional proteomics mass spectrometry data is collected using a data-dependent acquisition (DDA) protocol, in which each observed fragmentation spectrum nominally corresponds to a single, generating peptide sequence. This correspondence makes it relatively straightforward to frame the *de novo* sequencing problem as a spectrum-to-peptide translation task. In contrast, the alternative mass spectrometry protocol, known as data-independent acquisition (DIA) [12], yields data in which the signal associated with a single peptide sequence is distributed across multiple temporally adjacent mass spectra. Hence, rather than mapping from a single spectrum to a single peptide, in the DIA setting we are required to map from a series of spectra to a single peptide. Furthermore, in the DIA setting we must also take into account patterns in two different types of spectra: precursor (MS1) spectra as well as fragmentation (MS2) spectra.

Several existing methods are capable of carrying out *de novo* sequencing from DIA data. The first method involves extracting from the DIA data “pseudospectra” that resemble the MS2 spectra typically generated by a DDA protocol. The tool DIA-Umpire can be used for this extraction [13], and the resulting pseudospectra can then be fed into any existing DDA *de novo* sequencing tool [14, 15]. A significant drawback to this approach, however, is that a pseudospectrum is only created if a peptide generates detectable signal in the MS1 data. As a result, a significant portion of low abundance peptides are not detectable via this approach. Furthermore, due to the differences between the two protocols in how analytes are isolated and fragmented, DIA MS/MS spectra may look qualitatively different than spectra from the DDA experiments these models were trained on. The second method is a deep neural network, DeepNovo-DIA [16], that was specifically designed to operate on DIA data. It includes two convolutional neural networks and a long short-term memory network, which respectively capture 3D shapes in the MS2 spectra, correlations between MS1 and MS2 spectra, and peptide sequence patterns. A subsequent method, Transformer-DIA, adopts DeepNovo-DIA’s approach but swaps out the convolutional layers in the spectrum encoder with transformer self-attention layers [10].

In this work, we describe an alternative model, Cascadia, for *de novo* sequencing from DIA data. Formulating this model required solving two problems. First, we describe how to systematically extract small but complex data objects, called “augmented spectra,” that aim to capture all of the signal associated with a single peptide. Second, we describe how we modified the Casanovo transformer model [2] to take as input these augmented spectra. We train Cascadia from a large collection of annotated DIA data, and we demonstrate that the resulting model outperforms existing methods, including DeepNovo-DIA as well as Casanovo applied to DIA-Umpire pseudospectra. Tested on datasets collected on various instruments with different isolation window sizes, Cascadia consistently makes more than three times as many correct detections compared to other methods. Additionally, we verify variant peptides predicted by Cascadia using exome sequencing, and find that the highest confidence *de novo* peptides predicted by Cascadia show very high specificity.

## 2 Results

### 2.1 Cascadia operates on augmented DIA spectra

To perform *de novo* sequencing, Cascadia employs a transformer encoder-decoder architecture, similar to that of Casanovo [2] (Figure 1b). In this approach, the encoder first processes an observed spectrum, producing a latent representation of each peak. A transformer decoder layer than uses this spectrum representation, along with a learned embedding for each amino acid in the current peptide sequence, to autoregressively predict the next amino acid. However, generalizing this approach to work with DIA requires addressing two significant challenges.

**Figure 1:**
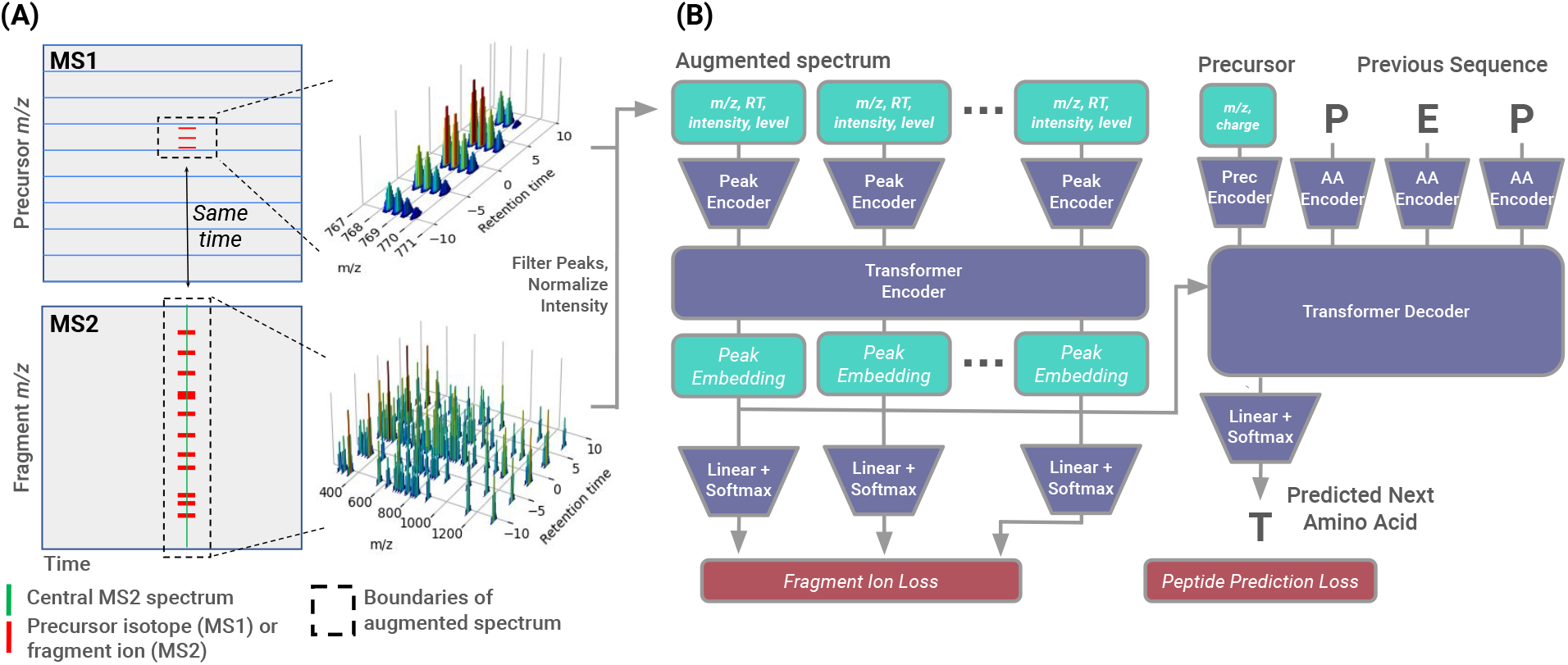
Cascadia schematic. **(a)** The figure illustrates an augmented DIA MS2 spectrum. Horizontal lines in the MS1 data correspond to the precursor *m/z* windows used in the DIA experiment. The matrix of MS2 data on the bottom consists of spectra collected from a single precursor *m/z* window, one spectrum per cycle. A selected MS2 spectrum (green vertical line) induces a series of fragmentation ladders over time as well as a corresponding series of isotope peaks over time (horizontal red lines). The augmented spectrum (dotted boxes) is designed to capture both of these patterns. **(b)** The figure gives an overview of the Cascadia workflow. Peaks in the augmented spectra are filtered, normalized, and encoded using *m/z*, retention time, intensity, and MS level embeddings. These peak representations are encoded by a series of transformer encoder layers and then passed, along with a sequence of learned embeddings for the previous amino acids in the peptide, to a transformer decoder layer to predict the next amino acid. A separate linear layer predicts, for each peak, whether that peak matches a *b* or *y* ion for the peptide.

The first challenge is that, in a DIA experiment, the signal corresponding to a single peptide is not captured in a single MS2 spectrum. This is because a DIA experiment measures every peptide species multiple times. The DIA setting involves isolating and co-fragmenting peptides in a pre-specified series of *m/z* ranges. The instrument fill time, the sizes of the pre-specified *m/z* ranges, and the chromatography settings are selected to ensure that each peptide is observed multiple times. For example, if the instrument carries out MS2 scans at a rate of 10 Hz, and if the DIA run is set up to iteratively sample a collection of 40 precursor windows, then the instrument will resample each precursor window once every ∼4 s. If a typical peptide species elutes from the liquid chromatography column for 15-30 s, then the instrument will observe that peptide 3–7 times. In addition, an MS1 scan is typically collected at the start of each cycle, and multiple successive MS1 scans contain characteristic isotope distributions for a given peptide. Thus, attempting to predict a peptide sequence from a single DIA MS2 spectrum would ignore valuable information from adjacent MS2 scans and from the corresponding MS1 scans.

One solution to this problem is to use a precursor detection algorithm to identify precursor isotope distributions that are observed in multiple successive MS1 scans and then construct an MS2 pseudospectrum for each identified MS1 feature. The pseudospectrum incorporates an accurate precursor *m/z* inferred from the peptide isotope distribution, and it aggregates information from multiple successive MS2 scans.

However, the pseudospectrum approach significantly reduces the potential sensitivity of a *de novo* method on DIA data because, in practice, many peptides can be detected in the MS2 signal despite not having a clear isotope distribution in the MS1 [17, 18]. In fact, in our data the proportion of peptides detected at a 1% FDR by the peptide-centric search tool DIA-NN which have a corresponding MS1 feature selected by DIA-Umpire is as low as 71.4% for our wide window test set collected on a Orbitrap Lumos and 47.2% for our narrow window test set collected on the Orbitrap Astral. In such cases, existing *de novo* sequencing methods will necessarily fail to detect the desired peptide, because no pseudospectrum for that peptide was ever constructed and passed to the model as input. This loss of sensitivity is why most database search tools for DIA analysis have moved away from using a pseudospectrum pre-processing step.

Cascadia takes an alternative approach to *de novo* sequencing of DIA data by expanding the inputs of the model to include all evidence for a given peptide, creating what we call an “augmented spectrum” (Figure 1a). To do so, we begin with a spectrum *S* that we hypothesize contains peaks associated with some generating peptide *p*, and we augment *S* with additional, nearby data that may also contain signals associated with *p*. The user specifies an augmentation width *w*, and the MS2 component of the augmented spectrum consists of spectrum *S* as well as the *w* spectra that precede it and the *w* spectra that follow it, within the same precursor window. In a similar fashion, the augmented spectrum captures isotope distributions in the MS1 data by incorporating MS1 peaks from the set of 2*w* + 1 MS1 scans that were captured just prior to each of the 2*w* + 1 MS2 scans. Furthermore, because the MS2 scans correspond to a specified precursor window, the augmented spectrum only contains MS1 peaks whose *m/z* values fall within this window (augmented by 2 *m/z* on either side in case the isotope distribution crosses the edge of the window).

In practical terms, this process of spectrum augmentation amounts to adding additional peaks to a given MS2 spectrum. A single MS2 spectrum *S* can be represented as a bag of *n* peaks (*m*_1_, *I*_1_), …, (*m*_*n*_, *I*_*n*_), where each peak has an associated *m/z* value *m*_*i*_ and intensity *I*_*i*_. In the DIA setting, we expand this peak representation to include two additional pieces of information: a Boolean *b*_*i*_ indicating whether the peak is from an MS1 (*b*_*i*_ = 0) or MS2 (*b*_*i*_ = 1) spectrum, and a time offset *t*_*i*_ indicating when the spectrum was measured relative to the central MS2 scan in the spectrum. Thus, we change our peak representations from (*m*_*i*_, *I*_*i*_) to (*m*_*i*_, *I*_*i*_, *t*_*i*_, *b*_*i*_), allowing the augmented version of spectrum *S* to be represented as a bag of 4-tuple peaks.

The second major challenge that we face in generalizing our transformer model to the DIA setting is the lack of accurate precursor information. During decoding, most DDA *de novo* sequencing methods condition the peptide prediction on a known mass and charge from the single precursor that was isolated for fragmentation. This information significantly narrows down the search space of potential peptide sequences. In contrast, in a DIA setting all analytes within the isolation window are isolated and co-fragmented, so the only available information about any given peptide is that its *m/z* value falls somewhere within a relatively wide range of values. DeepNovo-DIA solves this problem by inferring accurate precursor *m/z* values from the MS1 data, but at a significant cost in potential sensitivity.

In contrast, Cascadia skips the precursor detection step and instead passes the MS1 and MS2 peaks into the model directly. This approach gives Cascadia the flexibility to detect peptides without signal in the MS1 data while still capturing information from a precursor isotope distribution if one is present. At inference time, Cascadia processes all possible augmented spectra from a given mass spectrometry run, as opposed to just those with an identified precursor feature. Note that this strategy would not be feasible for the DeepNovo-DIA model, because extracting the peaks corresponding to theoretical fragment ions to pass as features to the Ion-CNN component of DeepNovo-DIA requires knowing the precursor mass *a priori*.

Another challenge we encounter when working with DIA data is that the spectra have a much lower signal-to-noise ratio and are are far more chimeric than DDA data, making the training signal for *de novo* sequencing much weaker. Unlike DeepNovo-DIA, which at each step only extracts potential fragment ions associated with the 20 possible amino acids, Cascadia considers all peaks in the mass spectrum at each step. Although this approach enables Cascadia to deal with missing fragment ion peaks, it also means that the model needs to learn the patterns of peptide fragmentation completely from scratch. Thus, we adopt two strategies to improve the learning process on noisier DIA data. First, we pre-train Cascadia’s spectrum encoder on high-quality DDA data from the MassIVE-KB repository, allowing it to learn the basic relationship between peptide sequences and fragmentation spectra before fine-tuning on the more complex DIA data. Second, we add an auxiliary classification loss term, where the model must predict whether each peak in the spectrum matches to a *b* or *y* fragment ion for the correct peptide. The motivation for this additional task, which is similar to the machine learning setup in the Spectralis *de novo* post-processor [19], is to provide a stronger training signal by explicitly indicating to the model which peaks in the mass spectrum are useful for predicting the correct peptide.

### 2.2 Cascadia outperforms state-of-the-art methods on wide-window DIA data

We hypothesized that Cascadia’s use of augmented spectra and a transformer architecture would yield substantially improved performance relative to DeepNovo-DIA. To test this hypothesis, we trained Cascadia on a collection of ∼5 million high-confidence peptide detections derived from 372 wide-window DIA mass spectrometry runs generated as part of the CPTAC consortium [20]. We generate training examples by searching the data with MSFragger-DIA and constructing one augmented spectrum for each peptide detected at a 1% FDR threshold. We then evaluate the model by performing inference on an entire held-out run.

One open question is how best to evaluate *de novo* sequencing tools on DIA data. In the DDA setting, precision and recall at the amino acid and peptide levels are commonly used as metrics to evaluate model performance [21]. This approach makes sense because the one-to-one correspondence between peptides and spectra means that a good *de novo* sequencing model is one that assigns the correct peptide label to as many spectra as possible. However, in the DIA setting, where evidence for each peptide is distributed across many spectra, this one-to-one correspondence is lost. To circumvent this problem, DeepNovo-DIA calculates precision and recall at the precursor level rather than the peptide level, which gives an indication of the fraction of the identified MS1 features that are successfully assigned a correct peptide sequence. However, results from this approach are confounded by the sensitivity of the precursor detection step and are not applicable to a method like Cascadia which makes predictions for all spectra at inference time.

As an alternative, we propose to evaluate DIA *de novo* sequencing methods at the peptide level by matching predicted peptides to a proteome database. To do so, we run each method on an entire unseen mass spectrometry run and take the maximum score assigned to each peptide sequence, after removing PTMs. At a given score threshold, we then obtain a lower bound on the peptide-level precision by using the protein database as a gold standard: each peptide prediction is marked as correct if it appears in the proteome for the species being analyzed. To reduce the likelihood of predictions randomly matching to the proteome, we restrict this analysis to peptides of length 8 or longer. This database matching approach allows us to directly compare the number of true discoveries made by each method at each estimated precision level. Furthermore, using a peptide-level procedure better reflects the primary metric of interest to most mass spectrometry practitioners, which is the total number of peptides detected in a given run, not the proportion of spectra or precursors that can be identified.

Using this evaluation protocol, we find that Cascadia discovers substantially more peptides across all estimated precision values than are found by DeepNovo-DIA (Figure 2a). For example, at a precision of 90%, Cascadia accurately predicts 2,371 distinct peptides, whereas DeepNovo-DIA only predicts 529. Furthermore, among the 2,371 high-precision Cascadia predictions, we find that 1,944 (82%) have a corresponding pseudospectrum. Hence, 427 of the additional high-confidence discoveries made by Cascadia are on analytes which lack a a clear MS1 signal and hence are not considered by DeepNovo-DIA.

**Figure 2:**
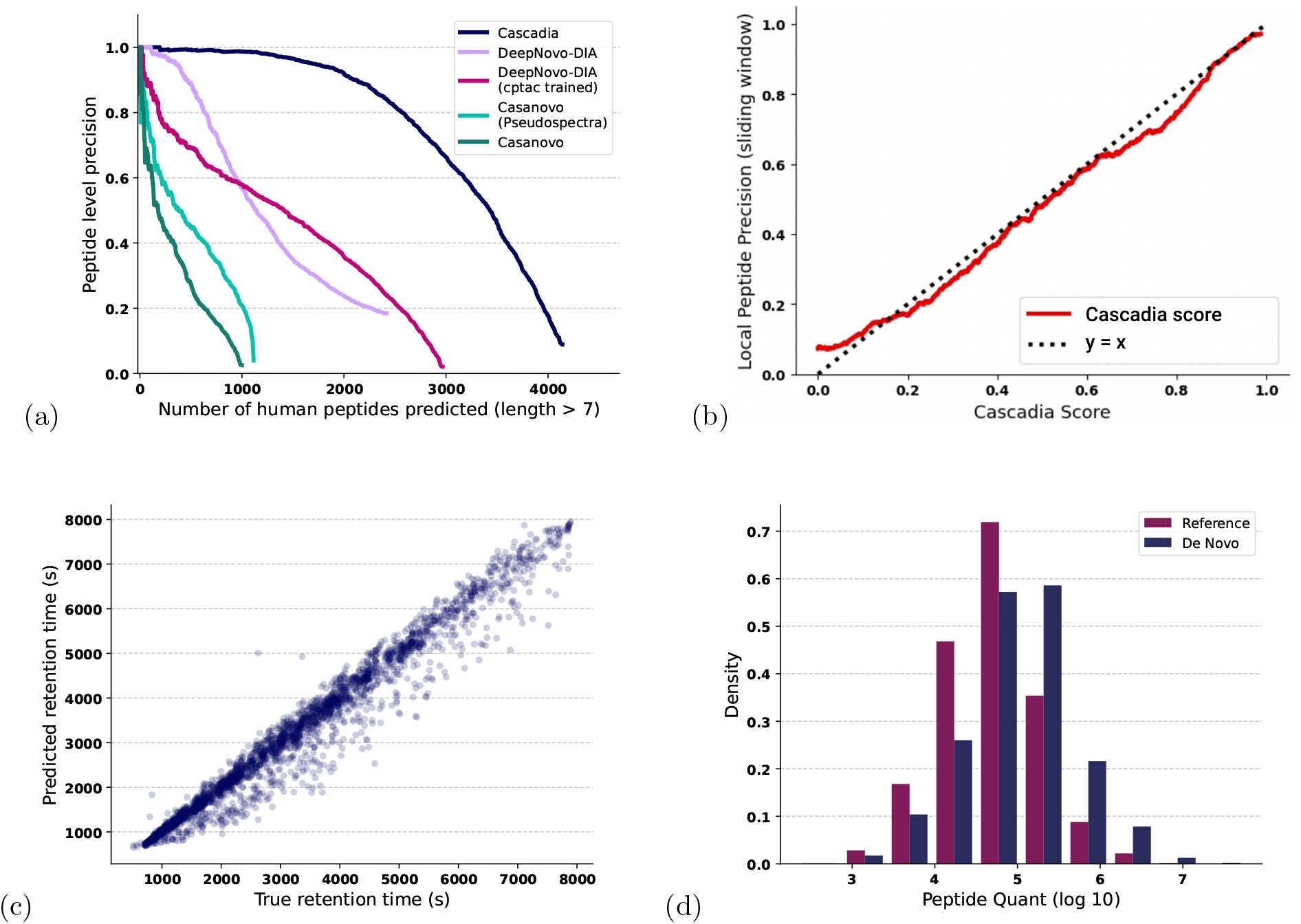
Comparison of de novo DIA tools on wide-window DIA data. **(a)** Precision-coverage curves showing the number of human peptides discovered by Cascadia, Casanovo, and DeepNovo-DIA on a single held out DIA mass spectrometry run from CPTAC. **(b)** A comparison between PSM level Cascadia confidence scores an local paptide precision, estimated using a sliding window of 500 PSMs. Cascadia scores appear to be relatively well calibrated across the range of confidences. **(c)** A scatter plot comparing the observed retention time for peptides predicted by Cascadia to the predicted retention time from AutoRT. **(d)** A histogram comparing peptide abundances between Cascadia *de novo* peptides and those that match to the reference.

To understand to what extent different training data contributes to this difference in performance between Cascadia and DeepNovo-DIA, we train a version of DeepNovo-DIA on the same CPTAC training set used by Cascadia. We find that although the resulting model predicts more total human peptides on the CPTAC test set, its confidence score is less well calibrated, resulting in fewer high-precision predictions (77 peptides at 90% precision).

Several publications have suggested that existing DDA *de novo* sequencing tools can be applied out-of-the-box to pseudospectra derived from DIA data [14, 15]. Thus, as a baseline, we applied Casanovo v4.1.0 to the pseudospectra as well as to the original MS2 spectra in our test set. As expected, using the original MS2 spectra yields very poor performance (29 peptides at 90% precision). Using DIA-Umpire extracted pseudospectra improves Casanovo performance somewhat (64 peptides at 90% precision) but is still far below the DIA-specific methods. These results are not surprising, as we would expect the wide-window DIA data used here to be significantly out-of-distribution for a model trained on DDA data.

To better understand the relative performance of Cascadia and DeepNovo-DIA, we also carried out a comparison using the plasma test set employed in the original DeepNovo-DIA paper [16]. To make a fair comparison, we provide Cascadia with augmented spectra that correspond to the precursor features used by DeepNovo-DIA. Note, however, that Cascadia does not have access to the inferred accurate precursor *m/z* values that are provided to DeepNovo-DIA. Despite this discrepancy, we observe that Cascadia again outperforms DeepNovo-DIA, as measured using spectrum-level precision and recall (Supplementary Figure S1). This result suggests that, in cases where there is a clear precursor isotope distribution, Cascadia is able to successfully infer it from the MS1 signal.

Next, to validate the set of *de novo* peptides from Cascadia that did not map to the reference, we compare the observed retention time for each detected peptide to the predicted retention time for that same peptide sequence from AutoRT [22] (Figure 2c). The 10,000 highest confidence detections from a standard database search with DIA-NN performed on the same run were used to fine-tune run-specific retention time predictions. Despite not taking as input the absolute retention time for each augmented spectrum, we see that the predicted and observed retention times are highly concordant for the majority of *de novo* peptides from Cascadia.

Finally, we compare the distributions of peptide abundances, as measured by DIA-NN, between the *de novo* peptides predicted by Cascadia and the database peptides detected by database search with DIA-NN (Figure 2d). Reassuringly, we see that the abundances for *de novo* peptides spans the same dynamic range as those detected by database search. Overall, the distribution for *de novo* peptides skews somewhat towards higher abundances, which likely reflects the greater level of evidence for a given peptide that is required to successfully *de novo* sequence compared with detecting that same peptide in a database search. In spite of this, Cascadia is still able to detect many analytes towards the low end of the abundance distribution.

### 2.3 Cascadia performs well on narrow window DIA data from the Orbitrap Astral

Advances in instrumentation continue to increase the resolution and acquisition speed of mass spectrometry experiments, allowing for the collection of DIA data at a faster throughput using smaller isolation windows. As instruments evolve, it is important that computational methods evolve with them. Accordingly, we evaluated the performance of Cascadia on DIA data generated by the recently released Orbitrap Astral. We train the model on 878,217 augmented spectra derived from 77 mouse plasma samples enriched for extracellular vescicles with the Mag-Net protocol [23]. We then evaluate the model on a held-out run generated from human plasma. For comparison, we also analyzed the human plasma data using DeepNovo-DIA and Casanovo.

The results of this evaluation again show that Cascadia provides a substantial boost in performance relative to DeepNovo-DIA (Figure 3a). At 90% precision, we observe an increase in the number of detected peptides from 2,137 to 6,224. On the other hand, naively applying Casanovo to DIA-Umpire pseudo-spectra performs much better on the Astral data than it does on the CPTAC data. This is likely because the isolation windows for the Astral dataset are much smaller than for the CPTAC data (4 *m/z* versus 16 *m/z*, respectively), and the resulting MS2 spectra therefore more closely resemble DDA spectra. Nonetheless, Cascadia still benefits from being able to see adjacent MS2 scans along with the full set of precursor peaks from the MS1.

**Figure 3:**
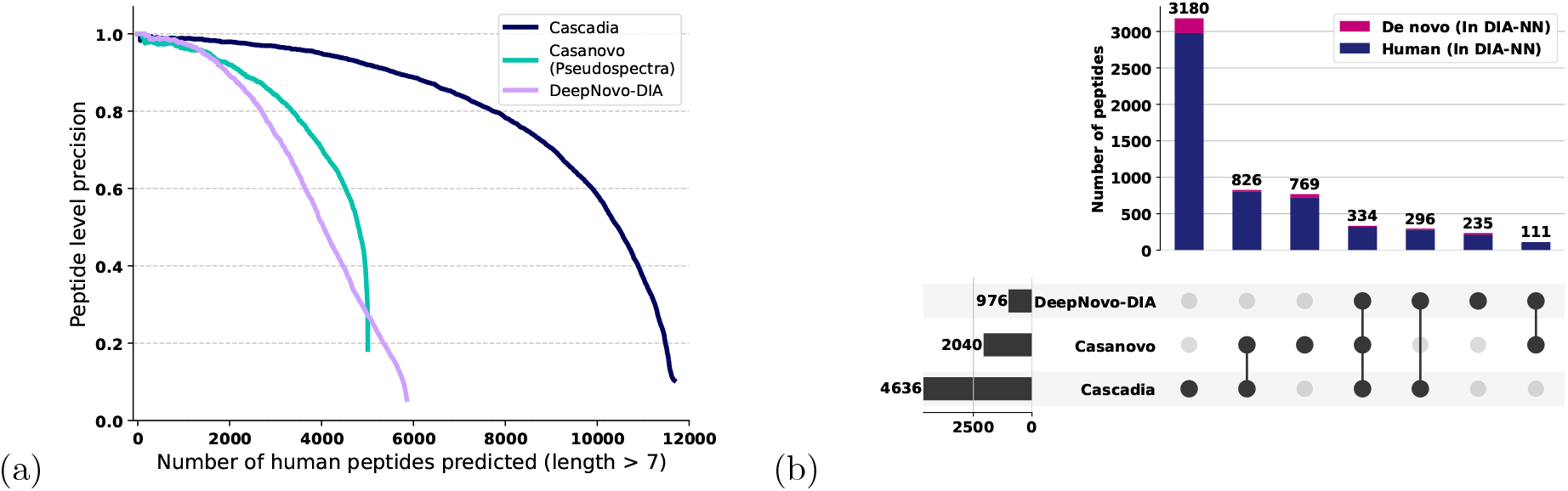
Comparison of *de novo* DIA tools on narrow window DIA data. **(a)** Precision-coverage curves showing the number of human peptides discovered by Cascadia, Casanovo, and DeepNovo-DIA on a single held-out Astral DIA mass spectrometry run. **(b)** An upset plot comparing the sets of high-confidence reference and *de novo* peptides discovered by Cascadia, Casanovo, and DeepNovo-DIA that are validated by a database search using DIA-NN. Predicted peptides are segregated into those that match exactly to the human proteome (red) or not (blue).

We also find that Cascadia scores are remarkably well calibrated as probabilities (Figure 2b), indicating that the model has learned an accurate representation of the conditional distribution of peptide sequences given observed spectra. This property makes it feasible to select a set of high-confidence discoveries by thresholding the Cascadia score. To better understand the sets of discoveries made by each of the three tools, we threshold predictions at the score threshold corresponding to 90% precision in Figure 3a for both Cascadia and DeepNovo-DIA, and we compare the predictions from each *de novo* sequencing algorithm to those found by the DIA database search tool DIA-NN [24]. To do so, we create a hybrid database containing the human reference proteome combined with the *de novo* predictions from both Cascadia and DeepNovo-DIA. In this experiment, we find that DIA-NN supports the majority of the Cascadia detections: of the 604 *de novo* peptides from Cascadia, DIA-NN also detects 316 (52%). On the other hand, of the 188 high confidence *de novo* peptides from DeepNovo-DIA, only 50 (26.6%) are supported by database search. Furthermore, comparing the sets of peptides (both *de novo* and reference) that are detected by each of the three methods and then validated by database search, we see that Cascadia both predicts the most total peptides and has the most overlap with the other two tools (Figure 3b).

### 2.4 Ablation experiments

To understand the effect of each of our model design decisions, we perform a series of four ablation experiments. Each of the first three models is pre-trained for the same number of epochs on the same MassIVE-KB DDA training set, and then all four are fine-tuned until validation loss converges on the CPTAC DIA dataset. The performance of each model is evaluated based on the number of human peptides correctly detected on a held-out CPTAC run.

The first two ablations involve removing components of the augmented spectrum: either eliminating MS1 peaks or eliminating peaks from adjacent MS2 scans. As expected, both of these modifications significantly hurt Cascadia’s performance, although removing flanking MS2 information appears to be much more harmful than removing MS1 information (Figure 4). We hypothesize that in many cases the model is able to infer the precursor information from just the fragment ions and the center of the isolation window, reducing the impact of not seeing MS1 peaks. On the other hand, we observed that, because the model without MS2 augmentation is unable to see the full elution profile, it frequently predicts the same peptide for many successive scans as opposed to only once at the center of the peak. Empirically, we observe that on average the MS2-deficient model predicts the same peptide in an average of 2.77 successive scans for a given precursor window, whereas the full Cascadia model predicts each peptide only 1.31 times in a row. The result of this redundancy is an overall decrease in the number of correctly predicted peptides.

**Figure 4:**
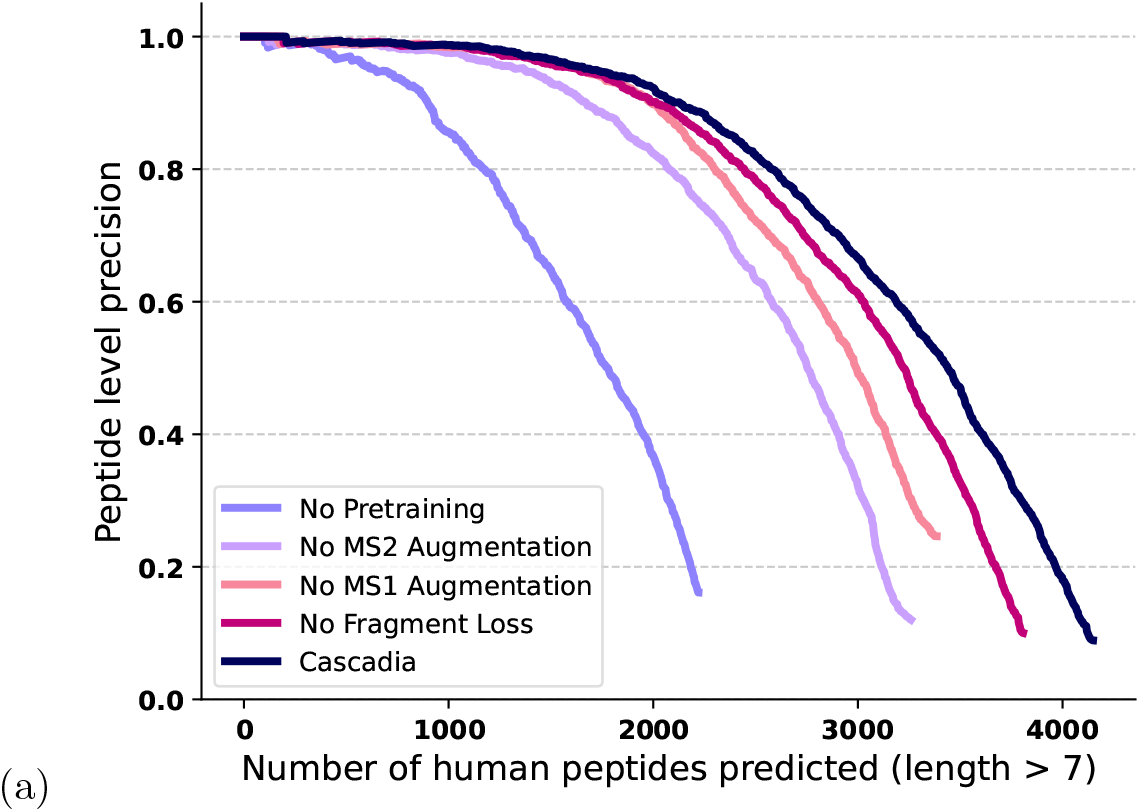
Ablation experiments. A series of precision-coverage curves showing how ablating various aspects of the Cascadia architecture affects performance.

The third ablation skips the DDA pre-training step. In this setting, the model training fails to converge on the much noisier DIA data and gives very poor performance.

The fourth ablation removes from Cascadia’s loss function the fragment peak prediction term, in which the model predicts which peaks correspond to b and y fragment ions. This change leads to a small drop in performance, but more strikingly increases the number of training iterations until convergence by 42%. We hypothesize that the additional task speeds up model training by providing a stronger counterfactual explanation for a given training example: rather than simply indicating the desired peptide sequence, the loss now also directly indicates which fragment ions might have been used to infer that sequence. Especially early in training, when the majority of model predictions are incorrect, this approach can help the model learn to deconvolve the relevant signal from the noisier and highly chimeric DIA spectra.

### 2.5 Cascadia discovers *de novo* coding variants from DIA data

To demonstrate Cascadia’s ability to discover *de novo* peptides, we test it in a setting where ground truth labels are available through an orthogonal sequencing modality. We generated three DIA runs on human skin samples derived from three different individuals. Targeted exome sequencing was then performed on 549 genes in these same individuals, yielding lists of 357, 368, and 595 ground truth single-nucleotide variants (SNVs) in each sample. Finally, we used Cascadia to *de novo* sequence the DIA data, and we used the Cascadia results to generate a list of candidate SNVs by searching for *de novo* peptides with exactly one mismatch to the human reference proteome and a confidence score above 90%.

The results of this analysis suggest that Cascadia can detect a subset of the SNVs, and that the Cascadia score accurately discriminates between correct from incorrect variants. In particular, of the 159 Cascadia predicted SNVs, 21 are confirmed by exome sequencing (9, 5, and 7 from the three samples, respectively). Furthermore, a receiver operating characteristic (ROC) curve, produced by ranking these SNVs by their Cascadia score and labeling them based on the exome sequencing, has an area of 0.92 (Figure 5a). Visualizing top-scoring Cascadia prediction both as an annotated spectrum (Figure 5b) and as an extracted ion chromatogram (Figure 5c), we see clear support for the variant peptide predicted by Cascadia, with numerous fragment ions supporting the variant sequence over the reference (20 additional examples in Supplementary Figures S2–S4). These observations provides evidence for the accuracy of the high-confidence *de novo* peptides predicted by Cascadia and suggest that *de novo* sequencing of DIA data may facilitate making additional discoveries that are missed by standard database search procedures.

**Figure 5:**
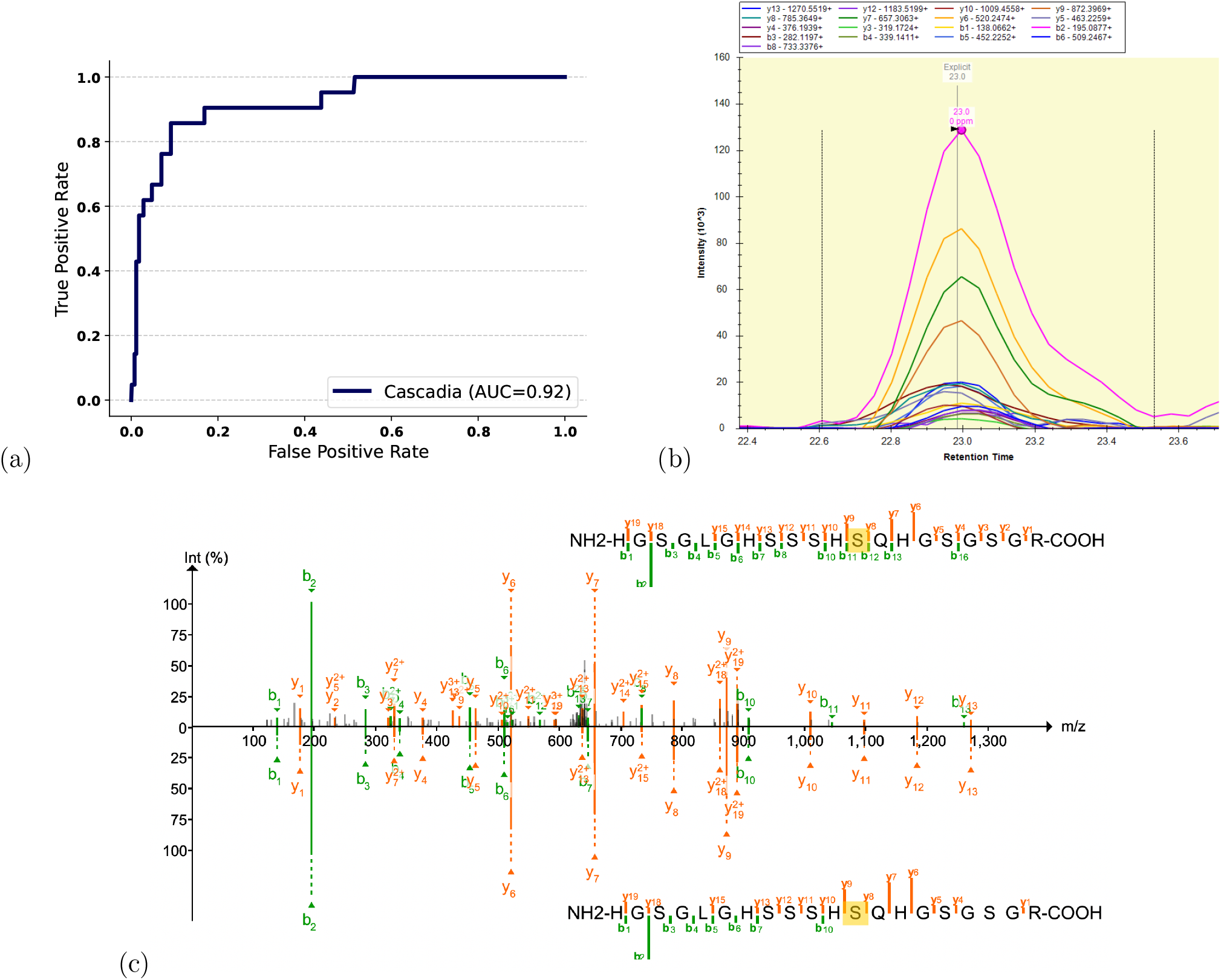
Variant prediction with Cascadia. **(a)** An ROC curve showing the ability of the Cascadia confidence score to separate correct and incorrect SNVs predicted with scores *>*90%. Cascadia was used to generate candidate SNVs for three individuals based on predicted peptides with exactly one mismatch to the reference proteome. The Cascadia confidence scores accurately discriminate between variants that are confirmed by exome sequencing and those that are not. **(b)** The Skyline extracted ion chromatogram for the most confident variant peptide predicted by Cascadia, HGSGLGHSSSHSQHGSGSGR, showing a clear peak. **(c)** A mirror plot comparing the MS2 spectrum at the peak retention time to the predicted spectral intensity from AlphaPeptDeep [25] for this same peptide. The site of the Gly-to-Ser variant is highlighted in yellow. Multiple fragment ions confirm the variant sequence.

We also note that Cascadia’s performance on these data, as measured by finding exact matches to the human reference proteome, is comparable to that on the CPTAC test set, indicating that Cascadia generalizes well to data from a different sample type, collected in a different lab, using a different isolation window size, and collected on a different instrument than the data it was trained on (Supplementary Figure S5).

### 2.6 Cascadia discovers *de novo* antibody sequences from human plasma

To further test Cascadia’s ability to discover *de novo* peptides from DIA experiments, we perform inference on a single Astral run from a human plasma sample enriched for extracellular vesicles [26]. Plotting the precision-coverage curve for this experiment, Cascadia’s performance initially looks significantly worse than for prior experiments, with a large number of high confidence peptides not present in the human reference (Figure 6b). However, searching these predictions against the reference we see that a large number of them match closely to immunoglobulin, with 3305 Cascadia peptides aligning to immunoglobulin reference sequences with 3 mismatches or fewer. Antibodies exhibit extremely high levels of sequence variation due to the processes of V(D)J recombination, junctional diversification, and somatic hypermutation [27]. Thus, we hypothesize that many of these non-matching predictions from Cascadia represent true antibody peptides rather than incorrect predictions.

**Figure 6:**
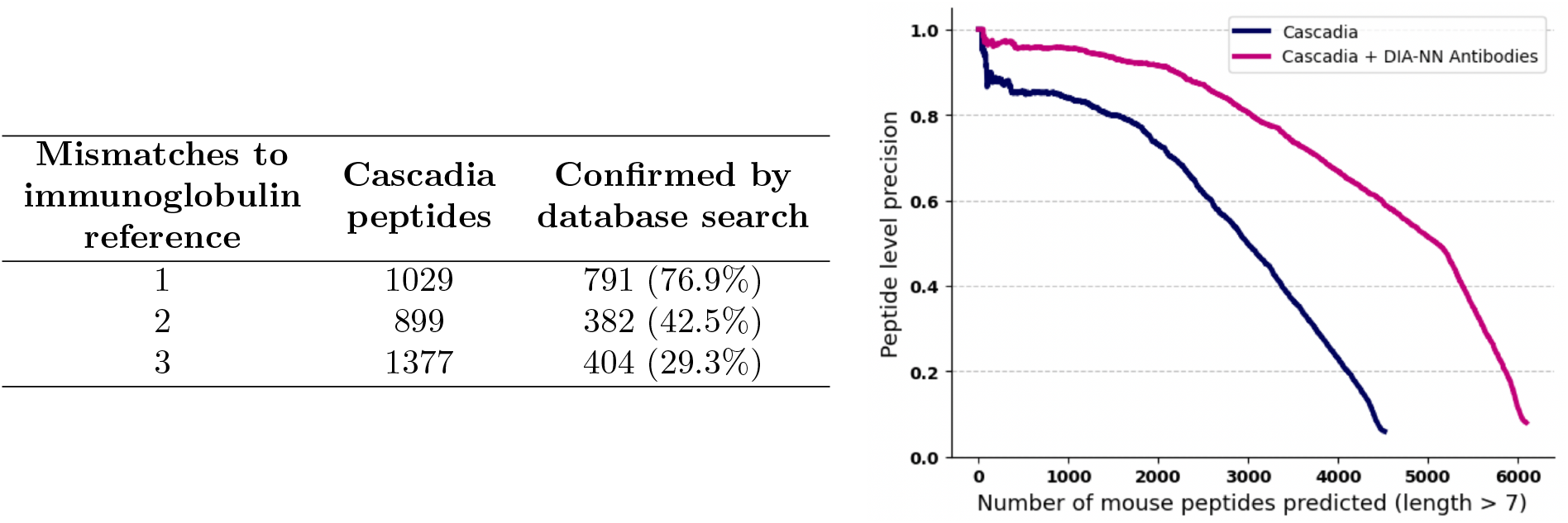
Discovering novel antibody peptides with Cascadia. **(a)** A table showing the number of candidate antibody peptides predicted by Cascadia that are validated by a database search with DIA-NN. **(b)** Precision-coverage curves showing the number of Cascadia detections before and after including the DIA-NN supported antibody sequences in the reference.

To test this hypothesis, we append the list of putative antibody peptides to the reference fasta and perform a traditional database search with DIA-NN. We reasoned that if DIA-NN also detects the same peptide with high confidence (*<*1% FDR), then this provides compelling evidence that the Cascadia predicted variant is correct. Of the 3305 candidate antibody peptides detected by Cascadia, 1577 (47.7%) are also detected by DIA-NN when included in the database (Figure 6a). Adding these validated peptides to the reference, the precision-coverage curve on this dataset improves significantly, showing similar performance to that seen on the other test sets (Figure 6b). Overall, this analysis demonstrates the ability of Cascadia to produce novel findings from existing DIA datasets and generate hypotheses that warrant further investigation.

## 3 Discussion

Cascadia is a transformer-based *de novo* sequencing model for DIA mass spectrometry data. Unlike existing DIA methods, Cascadia does not rely on an initial feature extraction step, instead operating on the raw MS/MS signal and extracting all relevant features directly. Benchmarking Cascadia on a diverse collection of data sets generated with different instruments and protocols, we find that Cascadia consistently offers state-of-the-art *de novo* sequencing performance on DIA data.

To evaluate *de novo* sequencing results for DIA, we adopt a new validation scheme that counts the number of distinct peptide sequences predicted by a given method that match to a reference proteome. In contrast to the spectrum-level metrics traditionally used to evaluate DDA *de novo* models, this peptide-level approach is in line with the way standard database search tools are evaluated for DIA data. Furthermore, it is worth noting that current approaches for evaluating *de novo* sequencing performance in the DDA setting are inherently biased, in the sense that they rely only on spectra that can be identified using a search engine. This approach makes the apparent *de novo* sequencing performance highly sensitive to the power of the underlying search: a less sensitive search is likely to include only high quality spectra which are easy to both identify and to *de novo* sequence. Additionally, the database search approach lends a bias in favor of models whose training data were generated with similar search parameters to the validation set, leading to unfair benchmarks when comparing different tools. Here, we avoid this bias by obtaining ground truth labels based on peptides mapping to the reference proteome rather than those from a database search. An important caveat to this approach is that it offers only a lower bound on the performance of each method. Thus, to both improve method evaluation and promote the adoption of *de novo* tools by the community, there is an ongoing need for rigorous false discovery rate control for *de novo* sequencing methods.

To further demonstrate Cascadia’s effectiveness, we test its ability to discover *de novo* peptides in two applications: variant prediction and detecting immunoglobulin variable region peptides from plasma. In both experiments, orthogonal evidence supports the validity of the most confident *de novo* discoveries made by Cascadia. Although the overall sensitivity of *de novo* sequencing is low compared to database search for DIA data, the specificity among top-ranking predictions is very high. This observation suggests that Cascadia is a potentially valuable tool for generating new discoveries in applications such as immunopeptidomics, forensic proteomics, and antibody sequencing. Furthermore, Cascadia may be a powerful complement to database search in settings where reference databases are either incomplete or intractably large, including proteomics experiments on understudied organisms, environmental and microbiome samples, and paleoproteomics.

We anticipate a number of future directions for work to further improve Cascadia on DIA data. First, Cascadia can be specifically fine-tuned for specific applications of interest, for example by training on a dataset of MHC bound peptides to improve immunoproteomics analysis [28] or non-tryptic data to ameliorate the tryptic bias of the model [29]. Furthermore, there are many successful ideas proposed by *de novo* methods in the DDA setting which may also be beneficial in the DIA setting, including the addition of additional auxiliary training tasks [4, 9], alternate decoding strategies [6, 15], and post-processing algorithms to refine predictions [19, 30]. Finally, future work may benefit from incorporating additional important features of the data which Cascadia currently does not include, such as the retention time, collision energy, and injection time for each scan.

The open source Cascadia software and associated model weights are available with an Apache license at https://github.com/Noble-Lab/cascadia. In addition to the commonly used mzTab fromat, Cascadia can also export predictions to the “.tss” format, which can be loaded into Skyline as a spectral library, allowing for the easy visualization of results [31]. As such, Cascadia can be directly incorporated into a standard DIA protomics workflow either in place of or alongside a traditional search engine.

## 4 Methods

### 4.1 Cascadia architechture

Before describing the model architecture for Cascadia, we begin with a brief overview of the Casanovo model that it builds upon. Casanovo uses a transformer encoder-decoder architecture to perform sequence-to-sequence translation, taking as input a sequence of peaks from an acquired MS2 spectrum, and predicting as output the sequence of amino acids for the generating peptide. This architecture gives Casanovo the flexibility to take in with spectra with variable numbers of peaks as input and predict peptides of variable length as output. Each peak is encoded with a positional *m/z* embedding and a learned intensity embedding, which are summed and passed as input into the transformer encoder. The self-attention mechanism in the encoder learns to capture the context between pairs of peaks in the spectrum, yielding a contextualized representation for the spectrum as a whole. This spectrum representation is then used as input to the transformer decoder. Decoding in Casanovo is autoregressive: at each step the model predicts the next amino acid in the sequence, conditioned on the previous amino acids, precursor information, and spectrum representation.. Decoding proceeds until a special stop token is predicted or the current peptide exceeds the precursor mass.

The primary difference between Casanovo and Cascadia lies in the spectrum encoder architecture. The first challenge is that, whereas the peaks in an MS2 spectrum can be naturally ordered by their *m/z* values, the peaks in an augmented spectrum occupy a two dimensional *m/z* -by-time space. This change is analogous to the difference between a transformer for language modeling, where words need to be embedded to capture their 1D position in the sentence, and vision transformers for images (ViTs), where embeddings must capture the 2D position of image patches. Thus, we take a similar approach to Dosovitskiy *et al*. [32] and encode peaks using a two-dimensional positional encoder which jointly assigns orthogonal embeddings to peaks along the *m/z* and time dimensions.

A second challenge is modifying the architecture to handle four-tuples rather than two-tuples. This is straightforward: the peak *m/z* and measurement time *t*_*i*_ are encoded using the 2D sinusoidal positional embedding function, and the remaining two values (*b*_*i*_, *I*_*i*_) are encoded via a learned linear layer. These three encoded vectors are concatenated, passed through a linear projection, and then input to the transformer.

The final change required to adapt the Casanovo architecture to DIA data is to remove the dependence on an accurate precursor. Unlike DDA data, where a precursor of known *m/z* and charge is detected in the MS1 and isolated for fragmentation, in DIA data all analytes within a relatively wide *m/z* window are subject to MS/MS. Accordingly, the only precursor mass information we provide to Cascadia is the center of the *m/z* isolation window for a given scan. In this way, we force Cascadia to infer accurate mass information, if any is available, directly from the MS1 input. This approach gives Cascadia the flexibility to detect peptides without signal in the MS1 while still capturing information from a precursor isotope distribution if one is present.

### 4.2 Training setup

Cascadia is trained in two stages. We start by pre-training Cascadia on a dataset of 28 million high-confidence DDA PSMs from MassIVE-KB. This training step allows the model to learn a useful initial representation for spectra and peptide sequences, including learning about patterns generated by peptide fragmentation on a clean dataset. During pre-training, we treat each DDA spectrum as an augmented spectrum with width *w* = 0. In addition to the single MS2 scan, we calculate a hypothetical isotope distribution for the peptide label using brainpy [33], and we include the top three most intense peaks in the MS1. Finally, we add uniform noise sampled from [−10, 10] to the precursor *m/z* to simulate the wide isolation windows encountered in DIA data.

Following DDA pre-training, we fine-tune Cascadia on DIA spectra. To ensure high quality training data, we train only on augmented spectra that contain evidence for at least 25% of the theoretical charge +1 and +2 b- and y-ions for the identified peptide. This setup ensures that we only train the model on spectra with a sufficient portion of the fragmentation ladder present to make *de novo* sequencing feasible. Although peptide-centric database search engines are often able to obtain high-confidence peptide detections based on a handful of transitions and an accurate precursor *m/z* value, many MS2 spectra do not provide enough evidence for successful *de novo* sequencing. By filtering for high quality spectra, we restrict model training to the setting where the desired task is possible, increasing the strength of the loss signal and speeding up convergence.

To further strengthen the training signal during fine-tuning on DIA data, we add an auxiliary fragment ion prediction task to the loss function. For each peak embedding obtained from the spectrum encoder, we use a linear layer followed by sigmoid activation to predict whether that peak corresponds to an expected b, y, or precursor ion for the labelled peptide. Binary cross-entropy loss is calculated for each peak, re-weighted to balance positive and negative classes (because the vast majority of observed peaks are not fragment ions), and added to the final *de novo* sequencing loss.

For both pre-training and fine-tuning, the data is split into train and validation sets containing 80% and 20% of the data, respectively, while ensuring that there are no shared peptides across datasets. Training proceeds until loss on the validation set does not decrease for 100,000 batches, which occurred after 2 epochs during pre-training and 7 epochs during fine-tuning. The peak learning rate is set to 1 × 10^−4^ for pretraining and 1 × 10^−5^ for fine tuning, with 100k batches of linear warm-up followed by a cosine shaped decay. A batch size of 16 is used for both pre-training and fine-tuning. Our final model consists of nine transformer layers in each of the encoder and decoder, an embedding size 512, and eight attention heads, yielding a total of 49 million trainable parameters. When not otherwise indicated, we use an augmentation width of 2, a maximum precursor charge of 4, and retain only the top 150 most intense peaks per individual (unaugmented) spectrum.

### 4.3 Testing setup

At test time, Cascadia is applied to every possible augmented spectrum from a given DIA run. This means that one augmented spectrum is constructed centered on each MS2 scan for each possible precursor charge allowed by the model (2, 3 and 4 by default). Decoding in Cascadia is identical to Casanovo: the model auto-regressively predicts a peptide sequence one amino acid at a time, at each step choosing the most likely next amino acid given the current predicted prefix until a special stop token is predicted. The default amino acid vocabulary for Cascadia includes the 20 canonical amino acids (with a fixed carbamidomethylation on cysteine), as well as four common post-translational modifications (oxidation, deamidation, acetylation, and carbamylation).

Similar to Casanovo, Cascadia includes a post-decoding filter that penalizes predicted peptides (by subtracting 1 from their score) whose masses lie outside a user-specified range of the precursor mass (Supplementary Figure S6). For this filter, Cascadia uses the entire isolation window (typically, 2–16 *m/z*).

### 4.4 Datasets

#### 4.4.1 MSKB DDA data

We pre-train Cascadia on the same dataset that was used to train Casanovo: 28M high quality DDA spectra from more than 1M distinct peptides derived from the MassIVE knowledge base (MassIVE-KB; v.2018-06-15) [34], a library of over 669 million HCD spectra compiled from 227 public human proteomics datasets. A database search identified ∼30 million of these spectra as high confidence PSMs by applying an extremely strict (∼ 0%) PSM-level FDR filter. These PSMs were randomly split at the peptide level into training, validation and test sets containing 28, 1 and 1 million PSMs, respectively.

#### 4.4.2 Wide-window DIA data

To fine-tune Cascadia for wide-window DIA data, we use a dataset of 4.8 million labeled augmented spectra derived from 373 runs in human kidney and pancreas, which are publicly available as part of the CPTAC consortium (PDC ID: PDC000341 and PDC000200). These data were collected on an Orbitrap Fusion Lumos instrument with a variable width isolation window, which was adjusted dynamically over the course of each run from a minimum width of 16 Th. The raw files were searched using MSFragger-DIA version 21.1 with the standard DIA Speclib Quant workflow and default settings [35]. For each detected peptide, we go back to the raw data to construct a corresponding augmented spectrum at the same *m/z* and retention time. To extract a list of precursor features for DeepNovo-DIA, we run DIA-Umpire within MSFragger [13], again using default parameters, and construct one test example for each extracted feature.

#### 4.4.3 Narrow-window DIA data

For our experiments with narrow-window DIA data, we use a training dataset of 878,217 labeled augmented spectra derived from 77 mouse plasma DIA mass spectrometry runs with 4 Th isolation window collected on the Orbitrap Astral (release pending). Samples were prepared with the Mag-Net protocol for enrichment of extracellular vesicles [23]. Peptide detections for training were generated using the DIA Speclib Quant workflow in MSFragger-DIA, and precursor features for DeepNovo-DIA and Casanovo were selected using DIA-Umpire.

As a test set, we use a single Astral run with 4 Th isolation window collected from HeLa cell lysates. For our experiments predicting *de novo* immunoglobulin peptides, we use another 4 Th Astral run analyzing human cell plasma enriched for extracellular vesicles. Both of these datasets are available at https://panoramaweb.org/AstralBenchmarking.url released with ProteomeXchange ID PXD042704 [26].

#### 4.4.4 DeepNovo-DIA data

We also evaluate Cascadia on the Plasma validation dataset used by DeepNovo-DIA, available through the MassIVE repository with accession MSV000082368. To ensure a direct comparison on the same set of metrics reported by DeepNovo-DIA, rather than running standard Cascadia inference on all augmented spectra in each dataset, we restrict our analysis to only the list of precursor features originally used by DeepNovo-DIA.

### 4.5 Evaluation metrics

To evaluate different *de novo* sequencing tools we use exact matches to the relevent proteome database to obtain a lower bound on model performance. For the human data in this study, we use the UniProt *Homo sapiens* reference proteome UP000005640, downloaded on 2/15/24. For the mouse experiments, we use the *Mus Musculus* reference UP000000589, downloaded on 4/29/24. To avoid matches arising purely by chance, we restrict analysis to only peptides of length eight or more. To plot a precision-coverage curve, we sort predictions by the confidence score assigned to each peptide. For a given score threshold *α*, we then calculate the precision as 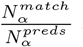 and coverage as 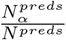, where 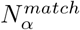 is the number of predicted peptides with score greater than *α* that match to the human proteome, 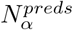 is the total number of peptides with score greater than *α*, and *N*^*preds*^ is the total number of peptides predicted by the model. With this approach, the evaluation does not depend on a database search engine as a gold standard. However, for consistency with prior work we also perform a comparison to DeepNovo-DIA using the same precursor level precision-coverage evaluation used by Tran *et al*..

### 4.6 Benchmarking

We benchmark Cascadia against the existing DIA *de novo* sequencing model DeepNovo-DIA, as well as to the performance of naively applying the DDA *de novo* sequencing model Casanovo directly to DIA data. Each method is run against an entire held-out mass spectrometry run and evaluated based on the estimated precision-coverage curve.

The DeepNovo-DIA code and model weights are downloaded directly from GitHub (https://github.com/nh2tran/DeepNovo-DIA). Because the feature selection algorithm originally used in DeepNovo-DIA is not publicly available, we use DIA-Umpire to generate a list of precursor features for inference. For evaluation on the the Plasma dataset that was originally used for benchmarking by Tran *et al*., we directly use the model predictions published by the authors in the MassIVE repository MSV000082368.

We also compare to the simple baseline of treating the DIA data as if it were DDA, and using Casanovo for inference. For this analysis, we use Casanovo 4.1.0 with its pre-computed model weights, and we leave all parameters as default. To make predictions on DIA data with Casanovo, we test two different approaches. For the first, we run Casanovo on only the central MS2 scan for each augmented spectrum used as input to Cascadia, providing only the center of the isolation window as a precursor. For the second, we run Casanovo on the Pseudospectra constructed by DIA-Umpire for each of the DIA-Umpire precursor features considered by DeepNovo-DIA.

### 4.7 Variant peptide prediction

To demonstrate the ability of Cascadia to predict novel *de novo* peptide sequences, we evaluate the model’s ability to detect novel coding single nucleotide variants (SNVs) from human proteomics data. We use a paired dataset of three DIA runs and corresponding exome sequences collected from three different donors (release pending). Targeted exome sequencing was performed on a shortlist of genomic co-ordinates corresponding to 549 genes, selected based on the the set of high-abundance proteins detected from the samples by an initial mass spectrometry run. For each mass spectrometry run, we perform inference with Cascadia and filter predictions at a 90% confidence score threshold. We then search the predicted peptides against the human reference proteome using pepmap version 2.0.0 (https://github.com/wenbostar/pepmap), allowing for up to one mismatch per peptide. The set of alignments with exactly one mismatch to one of the 549 genes targeted by the exome sequencing yields a set of candidate SNVs from the model. For each candidate SNV, we refer back to the paired exome sequencing data to obtain a ground truth label for whether the proposed variant is present in the donor. We then sort our list of candidate SNVs by Cascadia score, and we use the ground truth labels to plot an ROC curve, indicating how well Cascadia’s confidence scores are able to separate correct and incorrect variant peptides.

## Data availability

The MassIVE-KB DDA data used for pre-training is available at https://noble.gs.washington.edu/~melih/mskb_casanovo_splits.zip The wide-window DIA data from the CPTAC consortium used for training Cascadia is available through Proteomic Data Commons with study identifiers PDC000341 and PDC000200. The original DIA test set used by DeepNovo-DIA is available on MassIVE repository with accession MSV000082368. The Astral mouse training data will be available on Panorama Web (release pending), and the HeLa and human plasma EV astral test sets are available at https://panoramaweb.org/AstralBenchmarking.url with ProteomeXchange ID PXD042704. The data used in our variant prediction experiments will be published on Panorama Web (release pending).

## Code availability

Cascadia’s source code and trained model weights are available under the Apache 2.0 license at https://github.com/Noble-Lab/cascadia.

## Acknowledgments

This work is supported by National Science Foundation award 2245300, and in part by the Intelligence Advanced Research Projects Activity (IARPA) TEI-REX and PROTEOS programs under contracts W911NF2220059 and 2018-18041000004, respectively. The views and conclusions contained should not be interpreted as necessarily representing the official policies, either expressed or implied, of ODNI, IARPA, ARO, or the U.S. Government.

