## Supplementary figures for "A transformer model for *de novo* sequencing of data-independent acquisition mass spectrometry data"

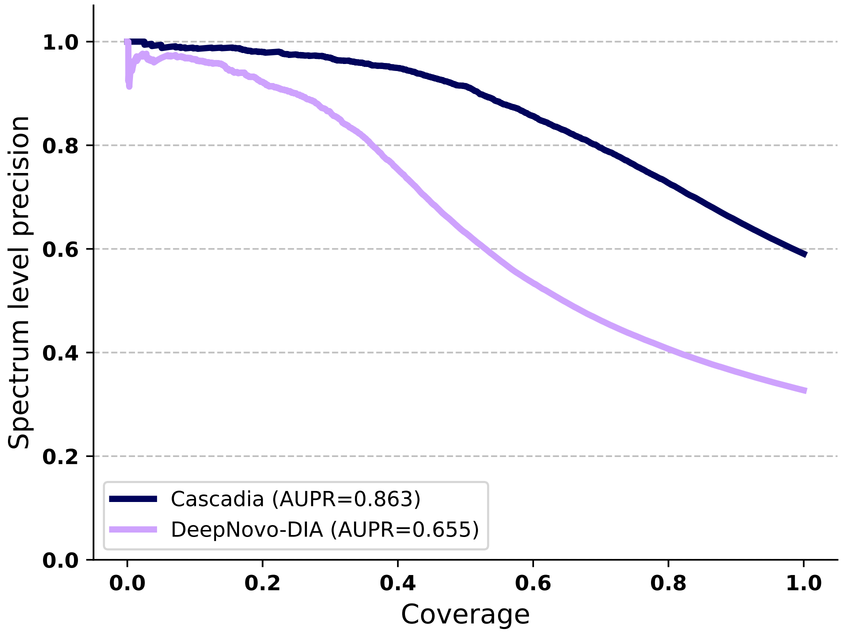


Figure S1: Spectrum level precision-coverage curves comparing the performance of Cascadia and DeepNovo-DIA on the same set of precursor features from the plasma test dataset used by Tran *et al.*

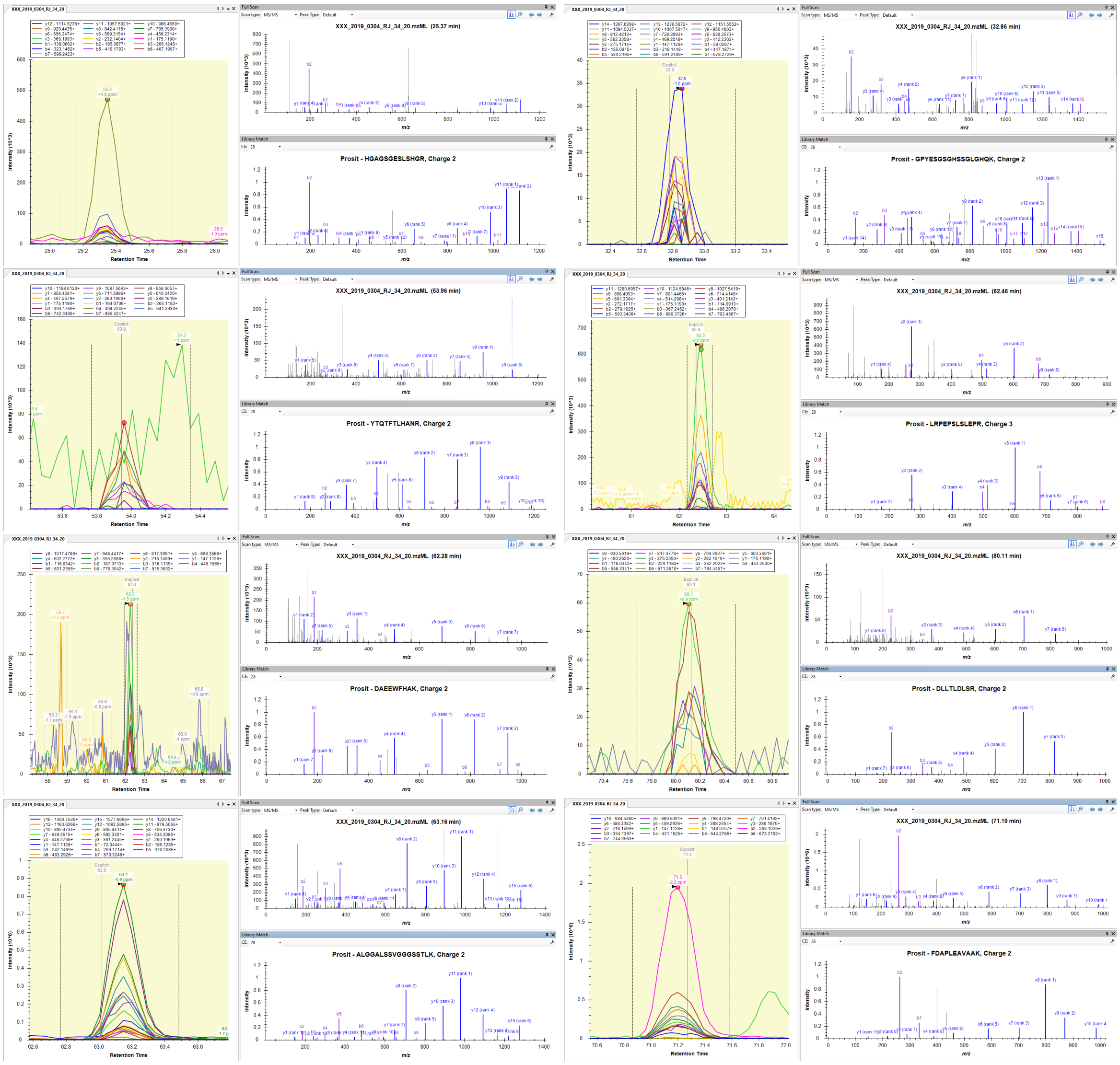

Figure S2: Skyline extracted ion chromatograms and a comparison between observed and Prosit predicted spectra for 8 variant peptides correctly predicted by Cascadia in donor 1.

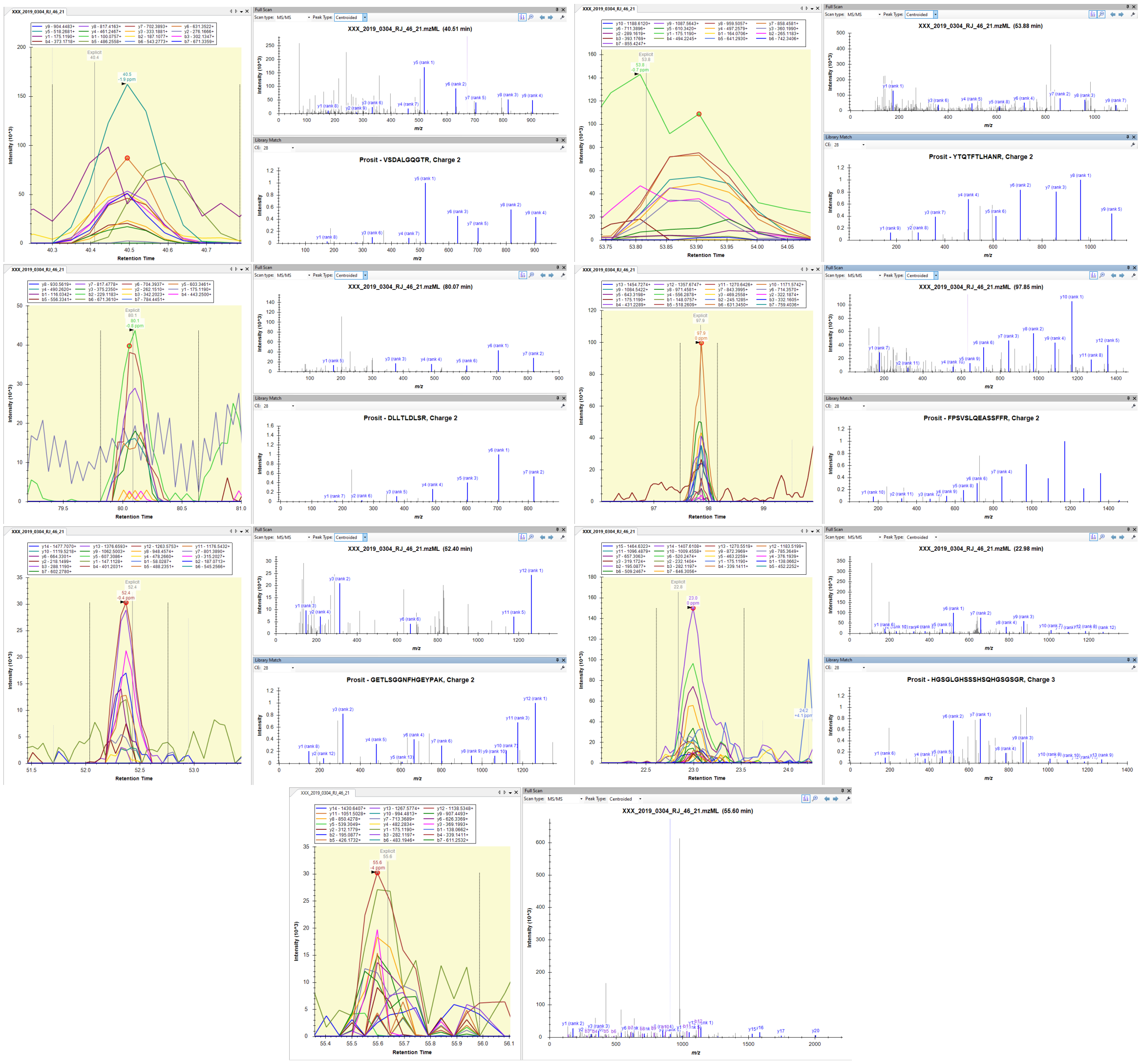

Figure S3: Skyline extracted ion chromatograms and a comparison between observed and Prosit predicted spectra for the 7 variant peptides correctly predicted by Cascadia in donor 2.

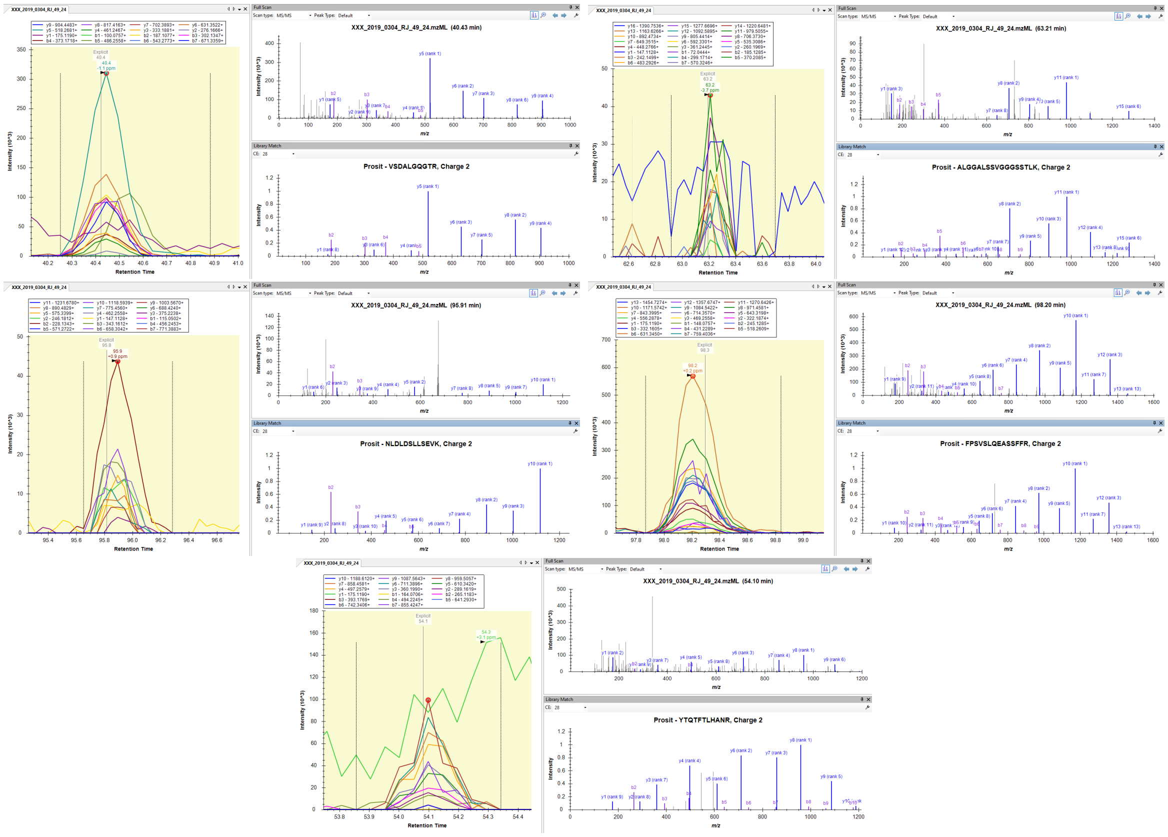

Figure S4: Skyline extracted ion chromatograms and a comparison between observed and Prosit predicted spectra for the 5 variant peptides correctly predicted by Cascadia in donor 3.

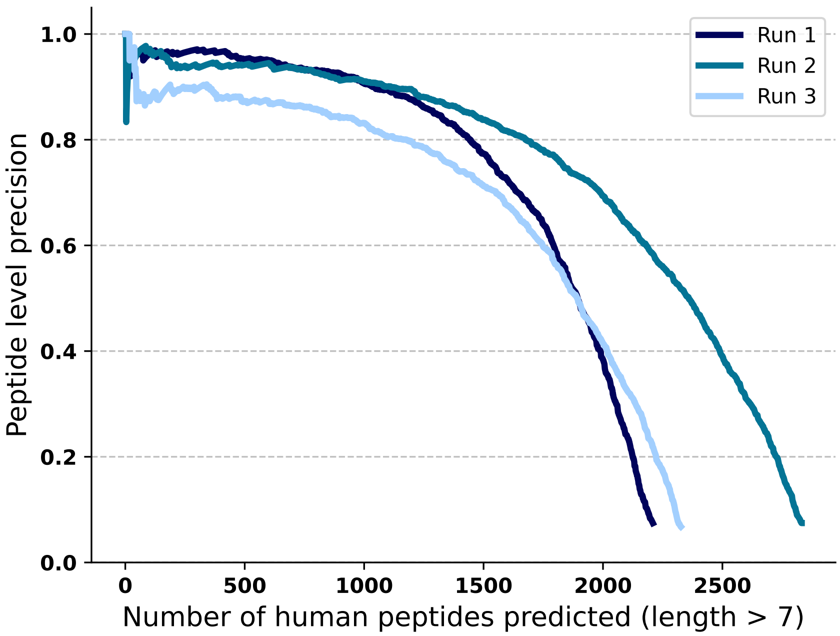


(a)

Figure S5: Precision-coverage curves showing the number of human peptides discovered by Cascadia on each of the three runs used in the variant analysis. Cascadia generalizes well to this data, despite it coming from a different instrument, samples type, and protocol than the training data.

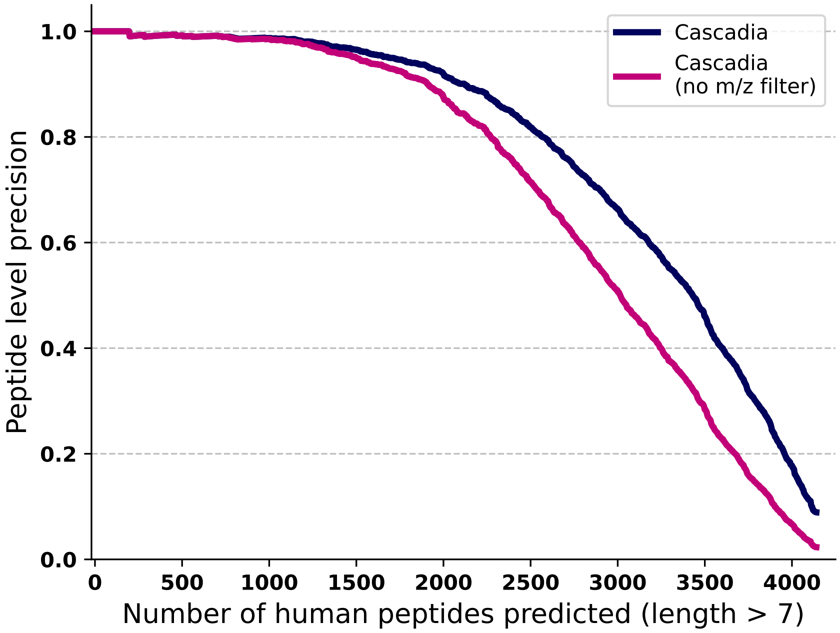


(a)

Figure S6: Precision-coverage curves showing the effect of the isolation window m/z filter on Cascadia predictions.
